# Energy-dependent protein folding: modeling how a protein folding machine may work

**DOI:** 10.1101/2020.09.01.277582

**Authors:** Harutyun K. Sahakyan, Karen B. Nazaryan, Arcady R. Mushegian, Irina N. Sorokina

## Abstract

Proteins fold robustly and reproducibly in vivo, but many cannot fold in vitro in isolation from cellular components. The pathways to proteins’ native conformations, either in vitro or in vivo, remain largely unknown. The slow progress in recapitulating protein folding pathways in silico may be an indication of the fundamental deficiencies in our understanding of folding as it occurs in nature. Here we consider the possibility that protein folding in living cells may not be driven solely by the decrease in Gibbs free energy and propose that protein folding in vivo should be modeled as an active energy-dependent process. The mechanism of action of such protein folding machine might include direct manipulation of the peptide backbone. To show the feasibility of a protein folding machine, we conducted molecular dynamics simulations that were augmented by the application of mechanical force to rotate the C-terminal amino acid while simultaneously limiting the N-terminal amino acid movements. Remarkably, the introduction of this simple manipulation of peptide backbones to the standard molecular dynamics simulation indeed facilitated the formation of native structures in five diverse alpha-helical peptides. Such effect may play a role during co-translational protein folding in vivo: considering the rotating motion of the tRNA 3’-end in the peptidyltransferase center of the ribosome, it is possible that this motion might introduce rotation to the nascent peptide and influence the peptide’s folding pathway in a way similar to what was observed in our simulations.

## Introduction

Once they are synthesized in a living cell, the majority of proteins rapidly attain their distinctive biologically active three-dimensional structures, called native conformations. These conformations are robustly achieved in vivo via a folding process that involves interactions of the folding chain with molecular chaperones and other maturation factors. The folding process often cannot be reproduced in vitro, in the absence of chaperones and other cellular components (1-5). However, some small proteins fold spontaneously in vitro in the absence of any other macromolecules (6).

What exactly happens during the folding of a linear polypeptide chain into a native conformation either in vivo or in vitro remains largely unknown. Despite decades of intense laboratory research, theory development and computer simulations, we still cannot recapitulate complete folding trajectories in silico, except for those of a few relatively short polypeptides (7). Knowledge of the intermediates in the folding pathways and the mechanisms that enable them is essential for determining the points of intervention at which folding and misfolding processes can be altered.

The painfully slow progress in our ability to fold in silico all but the shortest polypeptides could be due to the sheer complexity of the system: the number of possible conformations of a polypeptide chain, and the number of interactions between the atoms of all amino acid residues within the polypeptide itself and with the surrounding solvent, are so astronomically high that the existing computational power is not yet sufficient, and might never become sufficient, to capture the folding trajectories for longer proteins (8). It is also possible, however, that there are fundamental deficiencies in our understanding of folding as it occurs in nature, and progress in recapitulating protein folding pathways requires a more realistic physical model of folding than the one we have been relying upon.

The current dominant model of protein folding was prompted by early observations that some small proteins are able to fold in vitro into their native conformations spontaneously, in isolation from other proteins or cellular components (reviewed in (6)). These observations gave rise to the thermodynamic hypothesis of protein folding (6, 9), which in turn led to the development of the physical model that describes protein folding as a thermodynamically favorable, unassisted process. In a more recent, refined form, this model includes the description of a rugged funnel-shaped energy landscape, in which the various unfolded, unstructured conformations occupy the high-free-energy brim of the funnel (10-13). As the polypeptide chains fold, they sample conformations with progressively decreasing Gibbs free energy until they reach the native conformation, which is presumed to occupy the global thermodynamic minimum at the bottom of the funnel. The sampling of conformations during the folding process is assumed to occur via random thermal motions (14). The driving force of protein folding is assumed to be the decrease in free energy to the global minimum.

In summary, the current general physical model of protein folding describes a process that occurs in a closed system in the absence of external sources of energy. It assumes that folding starts from a random, unstructured conformation and proceeds unassisted, with no apparent requirement for the folding chain to interact with other proteins or macromolecular cellular components. This model describes an extremely artificial process that is only likely to occur in vitro and has little resemblance to what takes place during the folding of all proteins in the living cell.

In nature, folding of the majority of proteins occurs in the environment of a living cell, which is an open system with a constant flow of energy and shifting chemical composition. In a cell, a polypeptide starts folding while it is still being synthesized on a ribosome, where it occupies a tight space that allows it to adopt only a limited set of conformations. The nascent peptide emerges into a crowded, viscous environment outside of the ribosomal tunnel and interacts with multiple proteins, including chaperones, and with other cellular components, at all stages of folding. In the course of peptide synthesis and co-translational folding, a large amount of energy is released by GTP hydrolysis. This energy is not required for the formation of peptide bonds (15), but may be spent, at least partially, on various motions and adjustments of the ribosomal components, directly affecting the folding environment of the nascent peptide (16–18). It is difficult to escape the conclusion that protein folding in vivo must be described by a physical model that takes into account the interactions of a folding polypeptide chain with its complex dynamic cellular environment.

We have recently proposed that a more realistic physical model of protein folding might be built on the assumption that protein folding in vivo is an active, energy-dependent process. In this alternative model, proteins that are not able to fold spontaneously must rely on additional external forces to achieve native conformations (19). We hypothesized that the mechanism of action of such a protein folding machine might include direct mechanical manipulation of the peptide backbone by the concerted actions of the ribosome and chaperone complexes (20,21). During translation in the peptidyl transferase center of the ribosome, the 3’ terminus of the tRNA in the A-site swings by nearly 180 degrees in every elongation cycle (22,23). We hypothesized that this motion might lead to the rotation of the C-terminus of the nascent peptide. Simultaneously, the movements of the N-terminal regions of the nascent peptides may be restricted, first, by occlusions in the ribosome exit tunnel and then by steric capture mediated by the ribosome-associated “nascent chain welcoming committee", such as the trigger factor in bacteria and the nascent polypeptide-associated complex in archaea and eukaryotes (21). As a result the folding polypeptide may experience transient strained conformations with elevated free energy (19).

As the first step in exploring the feasibility of a protein folding machine capable of facilitating the attainment of native structure by mechanical manipulation of the peptide backbone, we performed molecular dynamics simulations augmented by application of torsion to the peptide backbones. During the simulations, the C-termini of various polypeptides were mechanically rotated either clockwise or counterclockwise, while the motions of their N-termini were restricted. We compared the trajectories of both types of simulations with the folding of the same peptides without the application of torque. In our experiments, directional rotation of the C-terminal amino acids with simultaneous limitation of the movements of the N-termini indeed facilitated the formation of native structures in five diverse alpha-helical peptides.

## Results

We performed atomistic molecular dynamics simulations to study peptide folding under conditions when, throughout the simulation, an external mechanical torque was applied to the C-terminal amino acid of a peptide and the motions of the N-terminal amino acid were restrained (Figure 1). We compared the folding trajectories of the peptide to which a mechanical force was applied to rotate the C-terminal amino acid in one of the two possible directions – either clockwise as in Figure 1, or counterclockwise – with the trajectories for the same peptide which was allowed to fold without any motion restriction or application of any mechanical force (referred to as “unassisted folding” below). As an additional control, we ran a fourth type of simulation, where motion restraints were applied to both ends of each peptide but the torque was omitted. The details of the simulations are described in the Methods section. Each of the four types of simulations were repeated three times, giving 12 simulations for each peptide.

**Figure 1.**
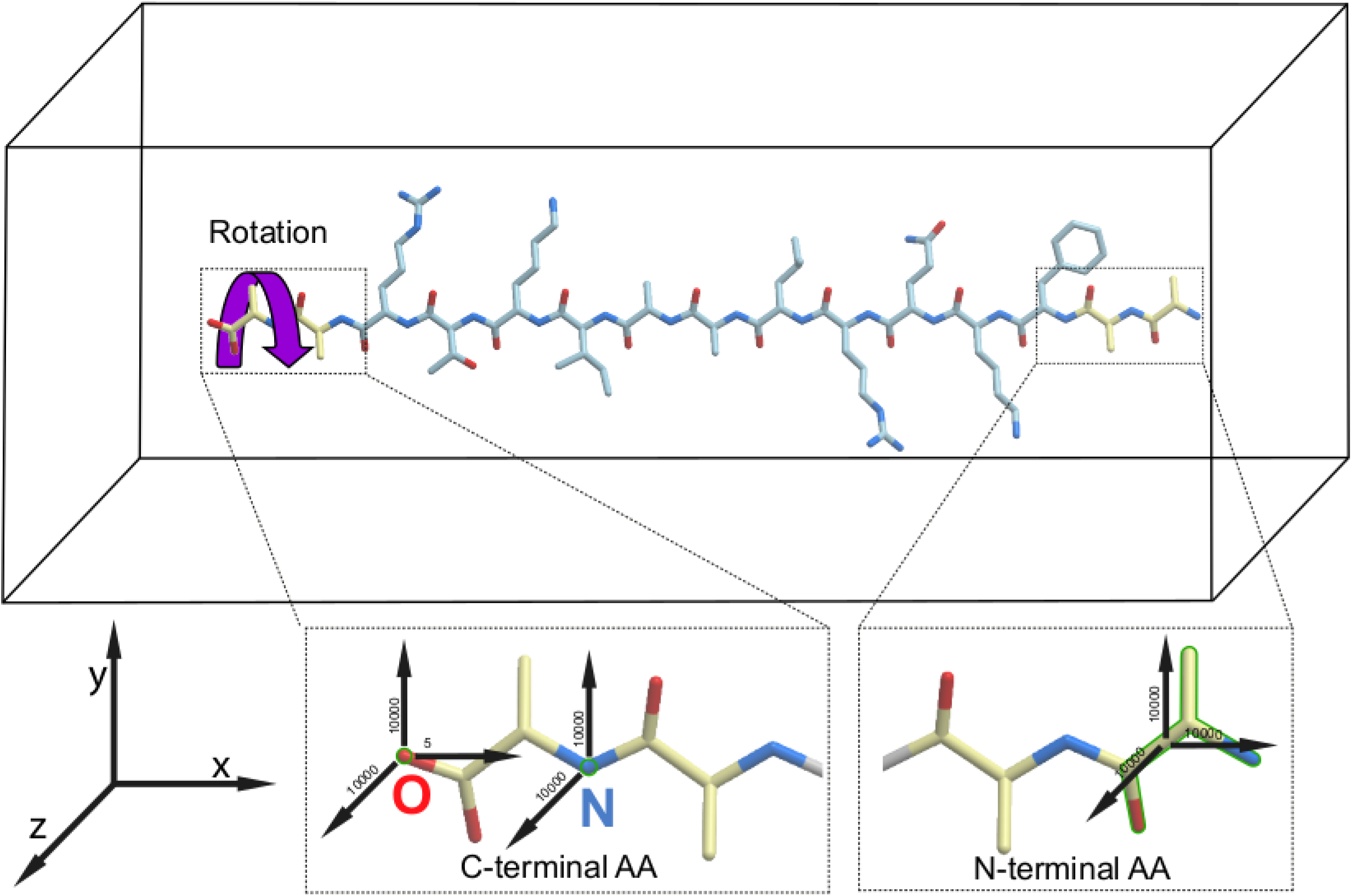
Schematic representation of the energy-dependent peptide folding protocol employed in this study. The force vectors applied to the C- and N-termini of a peptide in the simulation box are shown by black arrows. All force values are in kJ/mol*nm^2^. The purple curled arrow indicates the direction of the clockwise rotation of the peptides that resulted in the accelerated productive folding of all peptides to their helical conformations. The restrained groups are shown by green outline.

The experiments were run on five peptides that are known to adopt alpha-helical conformations in their folded form (Table 1). Two of these, P1 and P2, have been designed de novo, and the other three, P3-P5, are parts of naturally occurring proteins. The folding of the peptides was monitored by calculating the root mean square deviation (RMSD) distance of the peptide backbone from the native structure of the same fragment determined by X-ray crystallography (peptides P2-P5), or computed ab initio (peptide P1). The results of the simulations for each peptide when folded unassisted in the standard force field, and when an external torque force was added to the field, are presented in Table 2 and Figure 2. All folding trajectories are available at https://zenodo.org/record/4008419.

**Table 1.** Peptides used for the molecular dynamics simulations in this study.

| Peptide | Peptide description | Sequence | Length, amino acids | PDB ID |
| --- | --- | --- | --- | --- |
| P1 | Peptide Fs (Folded short), designed de novo | AAAA ( AAARA )<sub>3</sub> | 19 | n/a |
| P2 | First helix of the three-helix bundle, designed de novo | SWAEFKQRLAAIKTR | 15 | 2A3D |
| P3 | Fragment of the tetramerization domain of potassium channel Kv7.1 | HLNLMVRIKELQRRLDQSL | 19 | 6UZZ |
| P4 | Loop and third helix of the villin headpiece fragment HP35 | PLWLQQHLLKEKGLF | 15 | 2F4K |
| P5 | Fragment of the coiled-coil region of pyrin | KIQKQLEHLKKLRKSGEEQRS | 21 | 4CD4 |

**Table 2.** Peptide folding in the molecular dynamics simulations. The first number indicates the time (ns) spent before reaching the RMSD of 2 nm from the native conformation, and the second number indicates the duration of the experiment. 500/500 and 1500/1500 values indicate that folding was not observed in this simulation.

| Peptide | No rotation, no restraints | No rotation, restrained ends | Restrained ends, rotation clockwise | Restrained ends, rotation counterclockwise |
| --- | --- | --- | --- | --- |
| P1 | 500/500<br>500/500<br>500/500 | 500/500<br>500/500<br>500/500 | 22/100<br>18/100<br>29/100 | 500/500<br>500/500<br>500/500 |
| P2 | 1500/1500<br>1500/1500<br>1500/1500 | 1500/1500<br>1500/1500<br>1500/1500 | 160/200<br>140/200<br>105/200 | 1500/1500<br>1500/1500<br>1500/1500 |
| P3 | 1500/1500<br>1500/1500<br>1500/1500 | 1500/1500<br>1500/1500<br>1500/1500 | 105/150<br>95/150<br>125/150 | 1500/1500<br>1500/1500<br>1500/1500 |
| P4 | 1500/1500<br>545/1500<br>360/1500 | 250/1500<br>1500/1500<br>1500/1500 | 130/175<br>80/175<br>150/175 | 1500/1500<br>1500/1500<br>1500/1500 |
| P5 | 1500/1500<br>1500/1500<br>1500/1500 | 1500/1500<br>1500/1500<br>1500/1500 | 230/1500<br>380/1500<br>170/1500 | 1500/1500<br>1500/1500<br>1500/1500 |

**Figure 2.**
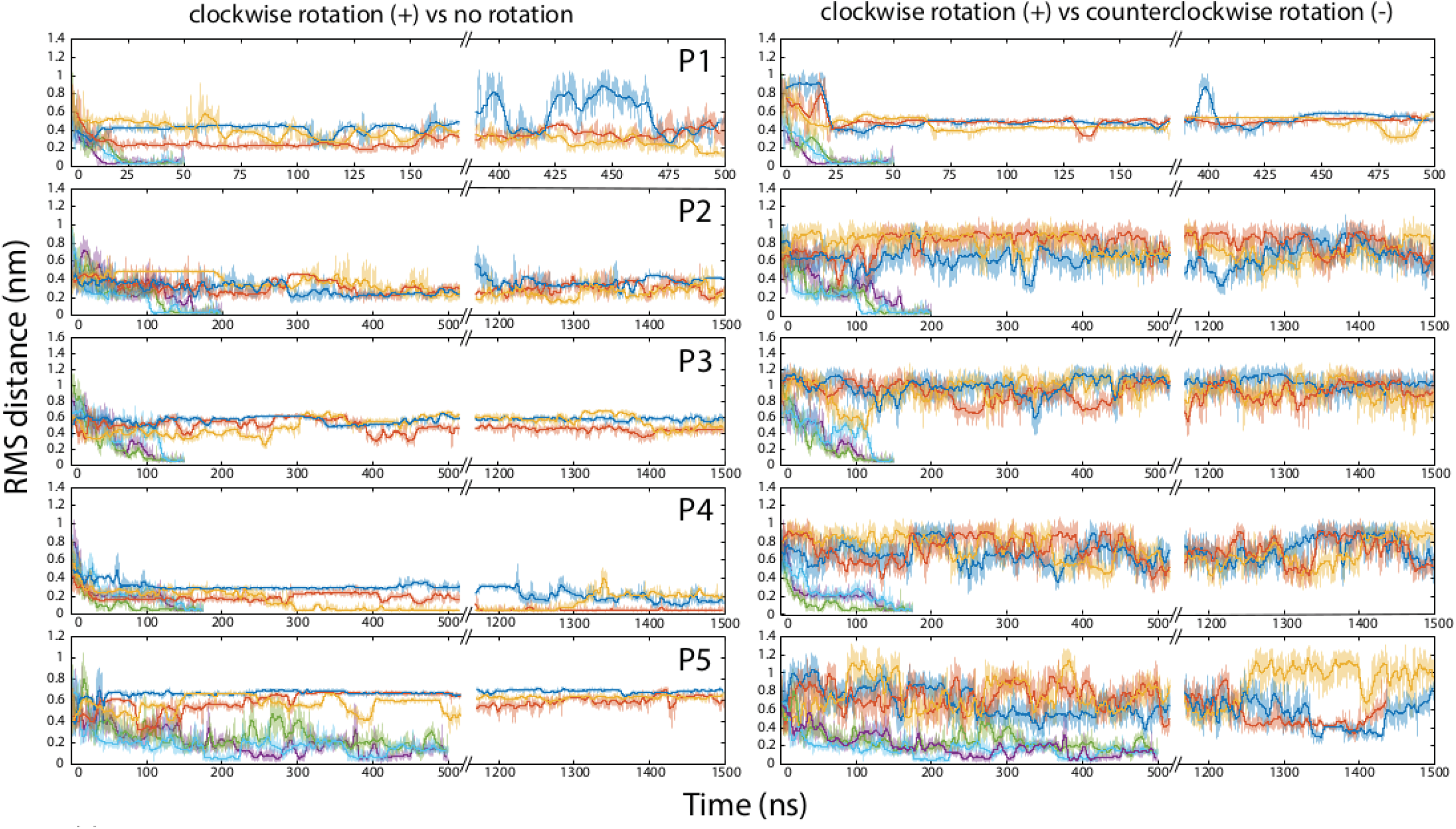
Folding of peptides in the force field with and without an augmentation by the application of external rotation forces to the polypeptide backbone. Each horizontal pane represents molecular dynamics simulations for one peptide, numbered P1 through P5 (Table 1). On the left side, top three curves (dark blue, orange, and yellow) indicate three independent runs for one peptide in the standard force field without externally applied backbone rotation, and the bottom three curves (purple, green, and light blue) indicate three runs in the presence of the clockwise rotational force. On the right side, the bottom three curves are the same as in the corresponding left pane (three runs in the presence of the clockwise rotational force), and the top three curves (dark blue, orange, and yellow) indicate three runs for the same peptide in the presence of the counterclockwise rotational force.

Within our simulation lengths, we observed the completion of unassisted folding into the native-like alpha-helical structure only in some runs for one peptide, P4, which represents the third helix and preceding loop in the villin headpiece domain HP35. Other peptides remained essentially unfolded throughout the 500-1500-nanosecond runs. The peptides also failed to fold when their ends were restricted in mobility but torque was not applied (Table 2). In contrast, when the external torsion force was applied to the C-termini of the peptides in the clockwise direction, as described in Methods and illustrated in Figure 1, peptides P1-P4 all folded into alpha-helical structures and were brought within 0.2 nm RMSD from their native structures in every run, typically within the first 100-200 ns of simulation. These peptides stayed in the native or nearly-native conformations for the remainder of the experiments. Peptide P5 was a special case; similarly to P1-P4, it adopted a compact conformation early in the experiments, but remained only partially folded for the duration of all runs (Figure 2).

For all five peptides, folding was observed when the rotation force was applied to the C-terminal amino acid in the clockwise direction (Figure 1). In contrast, the torque applied to the C-terminus counterclockwise with the same force constant did not facilitate folding of P1-P3 and P5, and may have inhibited folding of P4 (Figure 2).

## Discussion

To test the idea that inclusion of external forces can improve modeling of protein folding pathways in silico, we performed molecular dynamics simulations in which a standard force field was augmented by the application of external mechanical forces to the polypeptide backbone. We compared these simulations to control runs without any additional external forces. The directional rotation of the C-terminal amino acid with simultaneous restriction of the movements of the N-terminal amino acid facilitated the formation of native structures in five diverse alpha-helical peptides, confirming that such constraints can have significant consequences for folding dynamics. Strikingly, application of mechanical force accelerated the folding of P4, a fragment of an on-pathway folding intermediate of the well-studied villin headpiece domain HP35, which is one of the fastest-folding protein domains known (7,24,25). The several-fold increase in the rate of P4 folding that was achieved in our experiments seems to suggest that the postulated “physical limit of folding” of HP35 as a whole (24,26) could be overcome by a protein folding machine. The other four peptides in our experiments likewise attained their alpha-helical structure in the presence of the rotating force, but did not reach their native conformations when allowed to fold unassisted, even though we ran the control unassisted simulations for ∼10 times longer than the simulations that included the application of the external force (Table 2). Some of those peptides might take a very long time to reach their native conformations without application of an external force, whereas others might never fold unassisted, if their unfolded states are more stable than the folded conformations.

These results are in line with our protein folding machine hypothesis (19). They also support a hypothetical mechanism through which the machine would directly alter the conformations of proteins by applying mechanical force to the peptide backbone (20,21). The feasibility of such a mechanism, however, is dependent on whether the torsion applied at one point of a peptide would propagate through the rest of the peptide chain and affect the movements of the distal parts of the peptide. The peptide backbone is often viewed as a freely jointed chain, due to the 360-degrees rotation ability around the phi- and psi-bonds within each amino acid (27). If the peptides in our simulations were to behave as freely jointed chains, the rotation of a single amino acid at the end of the peptide would not have any appreciable effect on the motions of the rest of the peptide. However, if a mechanical torque were applied to a peptide while it was being folded in a viscous crowded environment (e.g., co-translationally in a living cell), we predicted that the free rotation of the phi- and psi-bonds in the peptide backbone would be hindered enough that escape from the forbidden sections of the Ramachandran plots would become difficult for many residues, and as a result, the entire peptide backbone may experience transient strained conformations. Although our simulation could not account for all the details of the protein folding environment in vivo, we were able to devise a set of conditions under which the peptide indeed did not behave as a freely jointed chain. When a force was applied to a single amino acid residue, and the motion of just one other residue at least 15 amino acids apart was restricted, the folding trajectory of the entire peptide was affected dramatically, leading to the rapid attainment of the native helical conformation. Some of the steric hindrances that make this rapid folding possible involve amino acid side chains, and therefore the effect might be sequence-specific. For example, glycine residues are more likely to experience the full 360-degree rotation around the phi- and psi-bonds, relieving the strain in the main chain; this might explain why P5, a peptide with an internal glycine, first acquired and then partially lost its folded conformation in our experiments (Figure 2).

It remains unclear whether our simulation captures the main features of the folding process as it occurs in nature. For example, one of the parameters that differs between our simulations and real co-translational protein folding process is their characteristic times. The rotation of the backbone in our system occurs at the submicrosecond time scale, whereas the addition of amino acids to the nascent peptide is much slower, on the order of subseconds (28-30). Molecular dynamics simulations have been known to model, at a fast scale, the essential parts of the molecular processes that are much slower when observed with bulk kinetics or single-molecule methods (17,25), but the effect of the rotation rate on the peptide folding trajectory remains to be investigated.

The key feature of the hypothetical mechanism of co-translational protein folding that we simulated is the directional rotation of the peptide backbone. As discussed above, the 3’ terminus of the tRNA in the A-site of the ribosome peptidyl transferase center turns by nearly 180 degrees in every translation elongation cycle. Only a 45-degree swing is necessary to achieve the proper stereochemistry of the peptide bond formation (31); the function of the remaining portion of the turn is unknown, and we have hypothesized that it may be needed to facilitate co-translational folding (20,21). It is notable, however, that the tRNA appears to turn in the counterclockwise direction when looking from the C-terminus of the nascent peptide (22,23). In contrast, folding of all peptides into the right-handed alpha-helices in our experiments took place only with clockwise rotation of the C-termini (Figures 1 and 2). It remains to be determined what, exactly, happens to the nascent peptide in the peptidyl transferase center and in the ribosome exit tunnel. The nascent peptide might be rotated counterclockwise (in the direction of the tRNA swing), or clockwise (as a result of a gear-like interaction with the tunnel walls), or might not be rotated at all but rearranged in a more complex way, being subject to pushing and pulling forces as well as interactions with the exit tunnel walls and other components of the ribosomal complex.

Regardless of whether the peptide torsion mechanism operates during co-translational folding on the ribosome in vivo, we demonstrate that it is possible to facilitate protein folding under conditions when an external mechanical force is applied to the peptide backbone. Importantly, we show that the peptide does not always behave as a freely jointed chain, opening the possibility that in vivo the peptide backbone can be manipulated into conformations that cannot be reached without assistance because they are either thermodynamically unstable or kinetically inaccessible. The results of our simulations thus demonstrate the feasibility of a protein folding machine. Some recently published results, including studies of the role of the exit tunnel in nascent chain folding (32-36) and of direct coupling between ATP hydrolysis and protein refolding by the chaperones of the HSP70 family (37-39), may be also interpreted as evidence of protein folding in vivo being an active process.

The notion of an active, energy-dependent protein folding mechanisms in vivo is better compatible with the current understanding of evolution than the generally accepted, standard thermodynamic hypothesis of protein folding. Although it is accepted that the ability of proteins to attain their native conformations must have evolved by natural selection of sequences that fold quickly and correctly (“evolution solved the protein folding problem” (40)), models of unassisted folding sidestep the fact that ribosomes and translation factors are among the oldest molecular machines shared by all extant cellular life (41), and were present during much of the evolution of proteins and of their folding pathways. The evolutionary optimization of the tempo and mode of protein folding, for at least 3.5 billion years of biological evolution, has taken place not in dilute solutions of isolated proteins, but in a dynamic environment of living cells with their constant flow of matter and energy. Thus, the ability of any present-day protein to fold in isolation and without assistance is likely to be either an incidental or derived property, not shared by most other proteins. Realistic computational modeling of protein folding must therefore take into account the presence of a multitude of external forces. Further studies should attempt to more closely recreate the conditions of protein folding in vivo.

## Materials and Methods

The initial stretched structures of peptides with four additional alanine residues, two at each end, were generated using ICM software (42). These alanines were attached as handles to which the rotation or restraint could be applied directly without affecting the sequence whose folding was investigated, and were not considered in the RMSD calculations. We aligned a peptide along the X-axis and solvated it in a dodecahedron box in the case of the simulations of unassisted folding and triclinic box in all other cases, with minimum distance of 1.5 nm between a peptide and the simulations box. Potassium and sodium ions were added to neutralize the charges in the system. The system was then minimized with the steepest descent algorithm, equilibrated for 100 ps in the NVT ensemble using V-rescale thermostat (43) for temperature coupling, and continued in the NPT ensembles for 1 ns using V-rescale thermostat and Berendsen barostat (44). After the equilibration, we kept temperature and pressure constant at 300 K and 1 bar respectively, using Nose−Hoover thermostat (45,46) and isotropic Parinello−Rahman barostat (47). The properties of simulation boxes for each peptide are summarized in Additional File 1.

For all simulations, we used the ff14SB force field (48) with the TIP3P water model (49) and ion parameters modified by Joung and Cheatham (50). Electrostatic interactions were calculated using particle-mesh Ewald (PME) summation (51) with a Fourier grid spacing of 0.135 nm. For non-bonded Coulomb and Lennard-Jones interactions, 1 nm cutoff was used. We constrained the hydrogen bonds with the LINCS algorithm (52) and used a 2-fs integration time step.

To exert an external mechanical torque to the C-termini of the peptides, we adopted the enforced rotation method, originally designed to study rearrangements during the rotation of a folded protein within the F1-ATPase assembly (53), implemented in the GROMACS molecular dynamics package. To this end, we restrained the positions of the O and N atoms of the C-terminal alanine to keep it aligned with the X-axis, about which the rotation was applied. The restraints with a force constant of 10000 kJ/mol*nm^2^ were applied only for the YZ-plane, so the C-terminal amino acid could move along the X-axis. In addition, we restrained the O atom of the C-terminal amino acid in the X direction with a force constant of 5 kJ/mol*nm^2^ and N and Cα atoms of the N-terminal alanine with a force constant 10000 kJ/mol*nm^2^ in all directions. The C-terminal amino acid was rotated using a flexible axis approach (Vflex2) with a reference rotation rate of 60 degrees/ps and a force constant of 1500 kJ/mol*nm^2^.

The GROMACS package version 2020.2 (54) was used for all simulations and trajectory analyses. The simulations were carried out on CUDA-enabled GPUs with Turing architecture, running Ubuntu Linux. For visualization of protein structures and trajectories, the programs ICM-Pro 3.9 (42) and VMD 1.9.3 (55) were used.

## Acknowledgments and Funding

We are grateful to many colleagues, most of all Dr. Yuri Wolf, for useful discussions. We thank Dr. Alexandra Mushegian for critical reading of this manuscript. INS is grateful to an anonymous angel investor for their support of Strenic, LLC. HKS and INS were funded by Strenic, LLC. HKS and KBN are supported by the National Academy of Sciences of the Republic of Armenia and Foundation Armenia (Switzerland). ARM is a Program Director at the National Science Foundation (NSF), the agency of the U.S. Government; his work on this project and stay at Clare Hall College, University of Cambridge, were supported by the NSF Long-Term Professional Development Program and Clare Hall Visiting Fellows Program, but the statements and opinions expressed here are made in a personal capacity and do not constitute an endorsement by the NSF or the Government of the United States.

