## Supplemental Table 1 for "Energy-dependent protein folding: modeling how a protein folding machine may work"

**Table S1. The details of simulation systems.** Simulations with restraints were performed in triclinic boxes and simulations without restraints were performed in dodecahedron boxes.

| System | Box volume | Total number of atoms in the box | Number of atoms within water molecules | Number of ions |
|---|---|---|---|---|
| **P1 (restrained)** | 101.11 $nm^3$ | 9826 | 3192 | 6/9 |
| **P1 (free)** | 351.01 $nm^3$ | 34456 | 11392 | 21/24 |
| **P2 (restrained)** | 121.26 $nm^3$ | 11787 | 3822 | 7/10 |
| **P2 (free)** | 186.68 $nm^3$ | 18128 | 5933 | 11/14 |
| **P3 (restrained)** | 131.49 $nm^3$ | 12759 | 4118 | 8/10 |
| **P3 (free)** | 157.44 $nm^3$ | 15248 | 4953 | 10/12 |
| **P4 (restrained)** | 109.71 $nm^3$ | 10642 | 3438 | 7/8 |
| **P4 (free)** | 148.58 $nm^3$ | 14381 | 4709 | 10/11 |
| **P5 (restrained)** | 164.43 $nm^3$ | 15965 | 5174 | 10/14 |
| **P5 (free)** | 251.53 $nm^3$ | 24378 | 7975 | 15/19 |
